# Intrinsic Network Connectivity Reflects the Cyclic Trajectory of Migraine Attacks

**DOI:** 10.1101/2021.06.30.450593

**Authors:** Anne Stankewitz, Enrico Schulz

## Abstract

**Background:** Episodic migraine is considered to be cyclic in nature, triggered by the hypothalamus. To assess the natural trajectory of intrinsic networks over an entire migraine cycle, we designed a longitudinal intra-individual study using functional magnetic resonance imaging (fMRI).

**Methods:** Intrinsic network connectivity was assessed for 12 migraineurs in 82 sessions including spontaneous, untriggered headache attacks and follow-up recordings towards the next attack.

**Results:** We found cyclic changes in the visual, auditory, and somatosensory networks, in limbic networks (e.g. thalamo-insular, parahippocampal), and in the salience network (anterior insula and dorsal anterior cingulate cortex). Connectivity changes also extended to further cortical networks, such as the central executive network, the default mode network, as well as subcortical networks.

Almost all of these network connectivity changes followed the trajectory of a linear increase over the pain-free interval that peaked immediately prior to the headache, and “dropped” to the baseline level during the headache. These network alterations are associated with a number of cortical functions that may explain the variety of ictal and pre-ictal physiological and psychological migraine symptoms.

**Conclusion:** Our results suggest that migraine disease is associated with widespread cyclic alterations of intrinsic networks that develop before the headache is initiated, i.e. during the interictal and premonitory phase. The increasing magnitude of connectivity within these networks towards the next attack may reflect an increasing effort to maintain network integrity.

## Introduction

Migraine is a cyclic disease that affects approximately 10% of the population. Recurring headache episodes are the most disabling symptom, but many patients also suffer from a variety of sensory and autonomous symptoms ^1^.

Migraine pathophysiology is complex and, thus far, not fully understood. Resting-state functional magnetic resonance imaging (rs-fMRI) has highlighted the presence of dysfunctional connectivities in migraineurs, particularly in somatosensory ^2^, dorsal attention ^3^, salience ^2^, executive control ^3^, and default mode networks ^3,4^.

Such a broad reorganisation of brain networks is thought to influence multisensory integration processes. In particular, alterations of limbic and sensory networks may lead to an increased sensory susceptibility that might make the migraineurs’ brain vulnerable to intrinsic and external factors that trigger migraine attacks. Despite the cyclic nature of the migraine disease, longitudinal studies are sparse ^5^. The majority of neuroimaging studies have used cross-sectional designs and found deviant cortical activity ^6,7^, seed-based connectivity ^8^, and resting-state network connectivity in migraineurs.

Here, we conducted an intra-individual rs-fMRI study to follow the trajectory of intrinsic cortical networks over the entire migraine cycle. Due to the clinical features of the disease and previous findings using neurophysiological and neuroimaging techniques ^9,10^, we hypothesised substantial alterations of sensory, limbic, and the salience networks during the migraine cycle.

## Materials and Methods

### Subjects

Twenty-two episodic migraine patients were included in the study, twelve of these data sets were complete and suitable for the statistical analysis (11 females and one male). Characteristics and clinical features are presented in Table 1.

**Table 1.** Intrinsic networks that showed activity changes over the migraine cycle.

|  | Increasing cycle | Decreasing cycle |
| --- | --- | --- |
| **Trajectory 1** | **visual (basal visual areas), auditory, somatosensory, central executive, salience, cerebellar, basal ganglia, pDMN, thalamic, frontal, temporal, sensory-motor, motor, limbic, insular, and cortically distributed (occipito-parietal and fusiform-parietal) networks** | **visual (higher order areas), cerebellar networks** |
| **Trajectory 2** | **-** | **limbic** |

Drop-outs from the initially-recruited subjects resulted from incidental MRI findings (n=2; acute mastoiditis and hydrocephalus), technical problems with the scanner hardware/ broken? head coil that lasted longer than four days (n=3) or software (n=1), illness of patients (n=2), or taking analgesic medication due to an acute painful event (n=1) during the scanning period. One patient decided to withdraw prematurely from the study. The exclusion of subjects was required due to the within-subject design, as the inclusion of incomplete data (predefined as a longer distance of 4 days between two scanning sessions) could have led to an inaccurate estimation of the subjects’ random effects in the statistical model. Moreover, an incomplete time series would miss mandatory data for the period immediately before the migraine attack.

Migraine patients were recruited via the interdisciplinary pain centre of the Klinikum rechts der Isar and online advertisements. Migraine diagnosis based upon the classification criterias of the International Headache Society ^11^ and was confirmed by a headache expert. The patients did not report any other neurological or psychiatric disorders, were not taking preventative medication for migraine for at least six months, but were allowed to take their regular acute migraine medication immediately (with a distance of 20 hours to the next scan) after the recording of the headache attack (non-steroidal anti-inflammatory drugs or triptans). All patients gave their written, informed consent. The study was conducted according to the Declaration of Helsinki and approved by the Ethics Committee of the Technische Universität München, Germany. All patients were remunerated for participation.

### Study design

Using a longitudinal, intra-individual study design, migraine patients were tested repeatedly over an entire migraine cycle (Figure 1). The imaging time series for each patient started with the recording of a spontaneous, untriggered and untreated headache attack within the first 4 hours after the beginning of the migraine headache. We only recorded headache attacks which were reported with an intensity of middle to strong, predefined as a minimum of “4” on a numerical rating scale with the endpoints zero (no pain) and 10 (extremely intense pain). Brain data were then recorded every 1-4 days at the same time of day (between 6 and 8 a.m.) until patients informed us by phone about the following headache attack (which was not scanned). The diagnosis of the second migraine attack was again confirmed by a headache expert. The time series was completed with the last attack-free recording.

**Figure 1.**
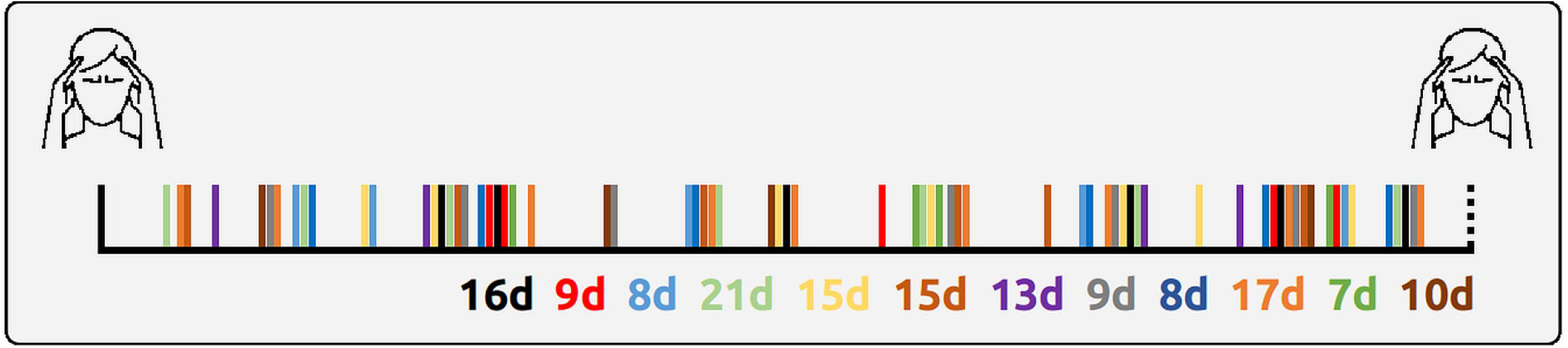
Time course of individual recordings. Solid vertical lines indicate recording days. The first and the last line represent the days of migraine attacks, the colourful lines in between represent the recordings within the migraine cycle relative to the attack days. Each patient has its own colour. The number of days between the first recorded migraine attack and the subsequent migraine attack indicates the different length of each patient’s migraine cycle (7 days for patient 11 and 21 days for patient 4).

We obtained data on the first day after the attack for all participants (12/12) and for the majority of the patients (9/12) on the day before or on the same day before the subsequent attack. All subjects had their final recording within 48h before the subsequent attack.

### Image acquisition

MRI data were collected on a 3 Tesla scanner (Ingenia, Philips, The Netherlands) using a 32-channel head coil. Patients were instructed to remain awake and relaxed with their eyes closed. For the 300 volumes of resting state data, we used the following parameters: TR = 2000 ms; time to echo (TE) = 30 ms; FOV = 192 × 192 mm^2^; flip angle = 90º; number of slices = 37; voxel size = 3 × 3 × 3 mm^3^ (0.29 mm gap). For image registration, a high resolution T1-weighted anatomical image was collected with: TR = 9000 ms, TE = 4 ms, flip angle = 8°, FOV = 240 × 240 × 170 mm^3^; number of slices = 170; voxel size = 1.0 × 1.0 × 1.0 mm^3^). Field maps were acquired in each session to control for B0-effects; 64 slices, TR = 960 ms, FOV = 192 × 192 mm^2^; voxel size = 2.0 × 2.0 × 2.0 mm^3^, 0.2 mm gap between slices. TE = 6 ms / 10.55 ms, flip angle 60°.

### Image preprocessing

The data were preprocessed with FSL. The Melodic toolbox was used to execute brain extraction, high-pass filtering with a frequency cutoff of 1/100 Hz, spatial registration to the MNI template, corrections for head motion during scanning, and - due to the potential contribution of smaller subcortical structures - a spatial smoothing (5 mm FWHM). The relatively small smoothing kernel was chosen to enable the detection of nucleic connectivity in subcortical regions. A distortion correction of the images was used based on field maps. The data were further semi-automatically cleaned of artefacts with ICA through Melodic. The number of components had been automatically estimated by Melodic and artefact-related components were removed from the data. Head movement during scanning did not exceed 2 mm or 2° in any direction. The movement during the migraine recording was within the range of the movement of the pre-ictal recordings (relative: 0.07±0.02 vs. 0.08±0.02; absolute: 0.2±0.7 vs 0.19±0.07).

### Statistical analyses

Following preprocessing, we ran a group ICA with temporally concatenated MNI-registered data of all 81 recordings using Melodic. Sixty-one networks (components) were determined by the ICA using automatic dimensionality estimation. Nineteen of these components were identified as artefacts by visual inspection (see Supplementary Material). Thus, we selected 42 resting-state networks. A dual regression analysis computed the individual maps for all 42 networks and each of the 81 sessions. We were running a voxel-wise statistics within the boundaries of the group network maps (as defined by Melodic) for the time series of 81 recordings. We aimed to investigate whether the trajectory of the time series is following two predefined time courses. In order to do so, we created two time vectors for each patient’s migraine cycle and encoded the day of the recording by assigning numbers between one and two. The numbers one or two were assigned to the measurements during the headache attack, depending on the following two possible trajectories of cortical processing (Fig. 2):

**Figure 2.**
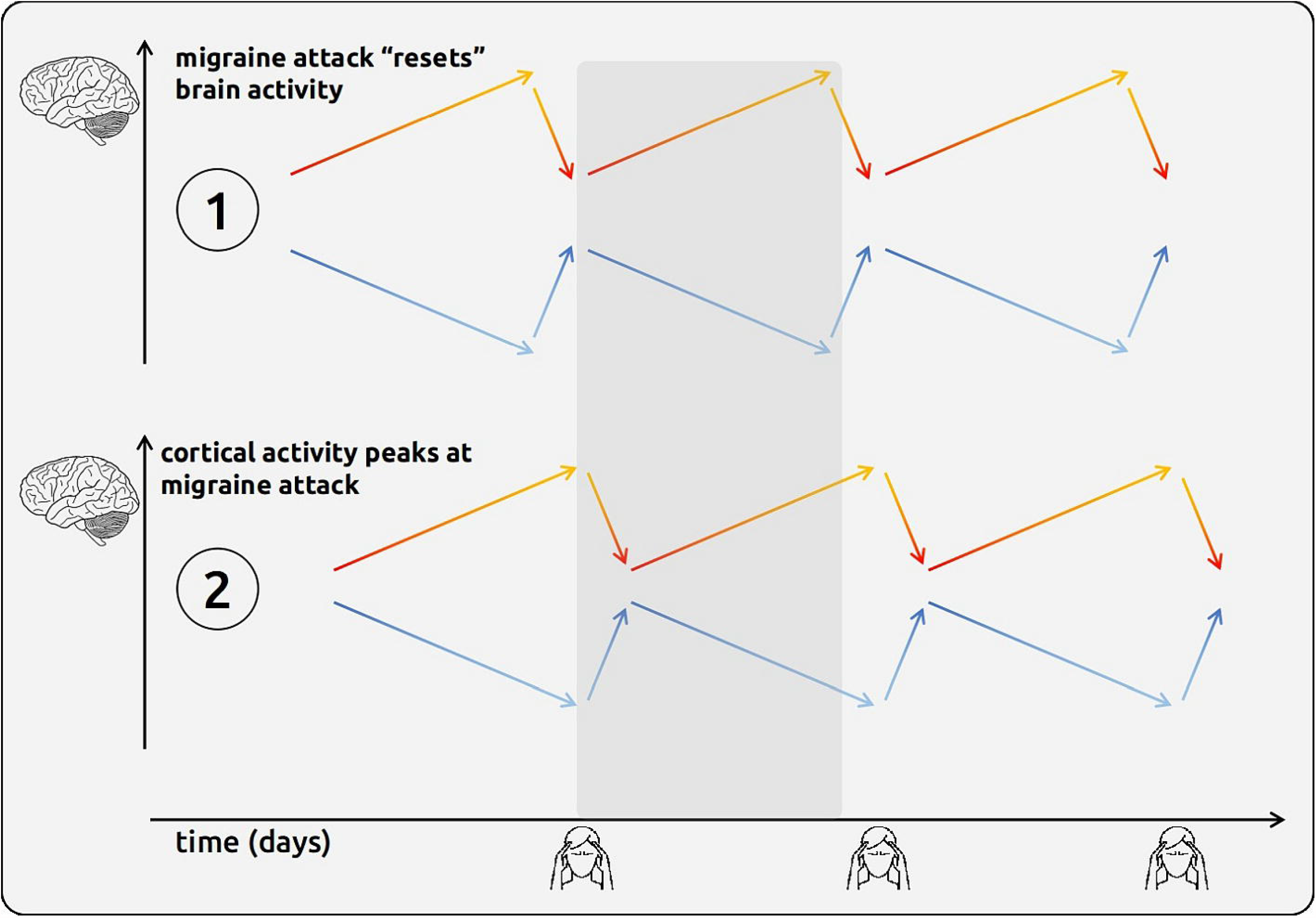
Time series of migraine-related resting-state maps. Two hypothetical time series of migraine-related cortical processes were modelled in the statistical analysis. In the first time course (upper part), the cortical processes drop during the headache attacks; the brain processes would be “reset” during attacks, then would resemble the processes on the day after the attacks. In the second time course (lower part), the cortical processes would reach their climax during the attacks and are similar to the days before attacks. These processes could be used as a biomarker for an impending migraine attack. **The figure is intended to illustrate the cyclic nature of migraine attacks and time-varying magnitude of two potential cortical processes; we recorded only one migraine cycle (grey area)**.

### Trajectory one (“reset mode”)

In this predefined model, we aimed to detect the resting-state connectivity that has its peak (or trough) just *before* the headache attack starts (during the premonitory phase) and then drops (or jumps) back to normal during the headache ^12,13^. From here, we model a linear increase (or decrease) over the migraine cycle to the next attack (Fig. 2, upper part). In this hypothetical time course, the magnitude of the cortical map on the first day after the attack would be similar to the brain connectivity during the attack. This trajectory can be interpreted as a cortical “reset” mechanism and is in line with neurophysiological and imaging studies suggesting altered cortical processes in migraineurs during the interictal interval that normalise just before or during headache attacks ^12,13^. For example, for five measurements over 10 days, the following vector is used to reflect trajectory one: 1 (=attack), 1.2, 1.4, 1.6, 1.8.

### Trajectory two (“pain mode”)

This model detects the cortical processes that have their peak (or trough) *during* the headache attack then drops (or jumps) back to normal the next day. From there, we assume a linear increase (or decrease) over the migraine cycle towards the next attack (Fig. 2, lower part). We hypothesised increased magnitude of brain connectivity in regions that contribute to the processing of migraine symptoms, e.g. pain, increased sensitivity to light, sound, and odours, and vegetative complaints. In this hypothetical time course, the brain connectivity on the day prior to the attack would be similar to the brain connectivity during the headache attack. Similar to the above-mentioned example with five measurements over 10 days, the following vector is used for trajectory two: 2 (=attack), 1.2, 1.4, 1.6, 1.8.

We explored the cyclic change of cortical rsfMRI connectivity over the migraine interval separately for each network. We computed voxel-wise linear mixed effects models (LME) in Matlab (Version R2018a, Mathworks, USA) within the z-masks of the network map as generated by Melodic. We related the time points within the migraine cycle to the session-specific cortical map of interest:

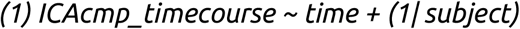

T-values of the fixed effect were computed voxel-wise as quotients between the beta estimates and the standard errors of the equation. The statistical results were corrected for multiple testing using the false discovery rate (FDR, p<0.05).

### Data availability

Inquiries for additional data are available upon reasonable request.

## Results

### Clinical characteristics

Twelve migraine patients were included in the study (mean age: 28; range: 21–40 years; 11 females). Migraine patients reported a mean disease duration of 13.6 years (range, 6-29 years). The attack frequency was recorded as the number of episodes per month and ranged between one and 10 per month. Nine patients reported migraines without aura. Laterality of headache pain was reported as either unilateral (five right-sided, three left-sided) or bilateral (four). Attack severity of the scanned headache was recorded on a numerical rating scale ranging from zero (no pain) to 10 (highest imaginable pain) and ranged between four and nine. The time points of the attacks were equally distributed across the menstrual cycles of the female patients. Characteristics and clinical features are presented in Table 1.

### Imaging results

We observed connectivity changes over the migraine cycle in 28 networks; 27 network changes apply to trajectory one and one network change applies to trajectory two. An overview of network changes depending on the trajectories is shown in Table 1; detailed information is given in Table 2. Maps of all networks are included in the Supplementary Material.

**Table 2a-c.** Cyclic changes of intrinsic networks.

**a. Trajectory 1** - linear *increasing* activity over the pain-free interval with a "drop" during the headache
| C | cluster sizes | t-values | MNI coordinates |  |  | Brain regions | Functional networks |
| --- | --- | --- | --- | --- | --- | --- | --- |
|  |  |  | x | y | z |  |  |
| 4 | 21 | 4.36 | -28 | -80 | -48 | Cerebellum | Attention |
|  | 18 | 4.6 | 40 | -60 | 42 | Lateral Occipital Cortex |  |
| 5 | 105 | 4.64 | 14 | -64 | 28 | Precuneus Cortex | Visual |
|  | 29 | 4.32 | -26 | -62 | 4 | Intracalcarine Cortex |  |
| 9 | 72 | 4.32 | 28 | -90 | 30 | Occipital Pole | Visual |
| 13 | 17 | 4.15 | 46 | -70 | -36 | Cerebellum | Cerebellar |
| 14 | 96 | 4.41 | -16 | -70 | 28 | Precuneus Cortex | Posterior DMN |
|  | 60 | 4.17 | 22 | -64 | 32 | Precuneus Cortex |  |
| 19 | 89 | 4.95 | -52 | -10 | 18 | Central Opercular Cortex | Temporal |
|  | 18 | 4.05 | 64 | -12 | 6 | Planum Temporale |  |
| 20 | 35 | 5.24 | -60 | -36 | 42 | Supramarginal Gyrus | Cortically distributed |
| 21 | 19 | 3.88 | 4 | -8 | 58 | Juxtapositional Lobule Cortex | Motor |
| 22 | 22 | 4.68 | -36 | 10 | -20 | Temporal Pole | Auditory |
| 25 | 20 | 4.22 | -2 | -30 | 26 | Posterior Cingulate Gyrus | Posterior DMN |
| 27 | 437 | 6.04 | -2 | -20 | 2 | Thalamus | Thalamic |
|  | 34 | 4.5 | 18 | -8 | -6 | Pallidum |  |
| 29 | 21 | 4.23 | -44 | -78 | 26 | Lateral Occipital Cortex | Cortically distributed |
|  | 18 | 4.48 | -62 | -60 | -6 | Middle Temporal Gyrus |  |
| 30 | 23 | 4.91 | -22 | 60 | -2 | Frontal Pole | Frontal |
| 32 | 31 | 4.24 | 24 | 0 | -8 | Putamen | Basal ganglia |
|  | 28 | 4.47 | -12 | 0 | 8 | Caudatus |  |
| 34 | 136 | 5.04 | 0 | 34 | 30 | Paracingulate Gyrus | Salience |
|  | 23 | 5.02 | 28 | 36 | 28 | Frontal Pole |  |
|  | 17 | 4.16 | -44 | -22 | 18 | Central Opercular Cortex |  |
|  | 16 | 4.71 | 40 | 12 | -6 | Insular Cortex |  |
| 35 | 29 | 4.53 | 56 | -34 | 4 | Superior Temporal Gyrus | Temporal |
|  | 20 | 4.7 | 30 | 20 | -20 | Frontal Orbital Cortex |  |
| 36 | 23 | 4.13 | -44 | -18 | 38 | Postcentral Gyrus | Sensory-motor |
| 37 | 39 | 4.46 | 48 | 26 | 0 | Frontal Orbital Cortex | Frontal |
|  | 18 | 4.01 | 52 | -8 | 46 | Precentral Gyrus |  |
|  | 18 | 5.15 | 18 | 20 | 68 | Superior Frontal Gyrus |  |
| 39 | 26 | 4.85 | 44 | -2 | -6 | Insular Cortex | Cerebellar |
| 43 | 126 | 5.44 | 36 | -44 | -28 | Temporal Occipital Fusiform Cortex | Cerebellar |
|  | 95 | 4.69 | -52 | -80 | 2 | Lateral Occipital Cortex |  |
|  | 50 | 4.33 | -16 | -60 | -12 | Lingual Gyrus |  |
|  | 20 | 4.37 | -50 | -68 | 20 | Lateral Occipital Cortex |  |
| 44 | 49 | 4.73 | -30 | -24 | 56 | Precentral Gyrus | Sensory-motor |
|  | 48 | 4.21 | -36 | -28 | 40 | Postcentral Gyrus |  |
|  | 22 | 4.58 | 12 | -52 | -16 | Cerebellum |  |
| 51 | 37 | 4.6 | 36 | 34 | 18 | Dorsolateral Prefrontal Cortex | Limbic |
| 58 | 152 | 5.52 | 32 | -6 | 46 | Precentral Gyrus | Sensory |
|  | 39 | 5.13 | 0 | -2 | 52 | Juxtapositional Lobule Cortex |  |

**b. Trajectory 1** - linear *decreasing* activity over the pain-free interval with a *peak* during the headache
| C | cluster sizes | t-values | MNI coordinates |  |  | Brain regions | Functional networks |
| --- | --- | --- | --- | --- | --- | --- | --- |
|  |  |  | x | y | z |  |  |
| 5 | 16 | 4.03 | 26 | -90 | 18 | Lateral Occipital Cortex | Visual |
| 43 | 100 | 6.1 | 22 | -90 | -18 | Occipital Fusiform Gyrus | Cerebellar |
|  | 21 | 4.6 | 0 | -52 | -22 | Cerebellum |  |

**c. Trajectory 2** - linear *decreasing* activity over the pain-free interval with a "*drop*" during the headache
| C | cluster size | t-values | MNI coordinates |  |  | Brain regions | Functional networks |
| --- | --- | --- | --- | --- | --- | --- | --- |
|  |  |  | x | y | z |  |  |
| 54 | 40 | 4.92 | -18 | -4 | -26 | Parahippocampal Gyrus | Limbic |

### Trajectory one

Changing connectivity predominantly applies to trajectory one, where 25 functional networks showed a linear increase of the magnitude of network connectivity over the interictal interval and a “drop” during the headache (Table 2a):

I. sensory: two visual networks (intra-calcarine cortex and occipital pole (=basal/ primary visual areas), lingual gyrus as well as precuneus), two somatosensory networks (post- and precentral gyrus, supplementary motor area, and cerebellum) and one auditory network (planum temporale and central operculum) networks;
II. two sensory-motor networks (precentral gyrus and supplementary motor area); III) central executive network (lateral occipital cortex and cerebellum);
III. salience network (dorsal anterior cingulate cortex, anterior insula, frontal pole, and central operculum);
IV. cortically distributed networks: two frontal networks (orbitofrontal cortex, frontal pole, and precentral gyrus), one temporo-supramarginal network (superior temporal cortex and orbitofrontal cortex), one occipito-lingual network (lateral occipital cortex and middle temporal gyrus) and one fusiform-parietal-frontal network (e.g. supramarginal gyrus and inferior frontal cortex);
V. two limbic networks (hippocampus and the dorsolateral prefrontal cortex (DLPFC);
VI. insular network (planum polare);
VII. thalamo-insular network;
VIII. basal ganglia network (putamen and thalamus);
IX. posterior cingulate-precuneus network (precuneus);
X. lingual network (posterior cingulate cortex, precuneus, supracalcarine cortex); XII) three cerebellar networks: intra-cerebellar (cerebellum), cerebello-insular (hippocampus) and cerebello-ponto-thalamic (temporal and lateral occipital cortices, cerebellum and lingual gyrus);
XI. posterior default mode network (DMN) (posterior cingulate cortex).

In two of these 25 networks, we also found brain regions that showed the inverse behaviour, namely a linear decrease of the magnitude of network connectivity over the pain-free interval with peak magnitude during the headache (Table 2b):

I. visual network (lateral occipital cortex (= higher order visual area) and the posterior cingulate cortex); and
II. cerebello-ponto-thalamic network (occipital fusiform cortex and cerebellum).

Figure 3 shows exemplary results in the visual (Fig. 3a), thalamo-insular (Fig. 3b), and the salience networks (Fig. 3c). All statistical tests were thresholded at p<0.05 (FDR corrected).

**Figure 3.**
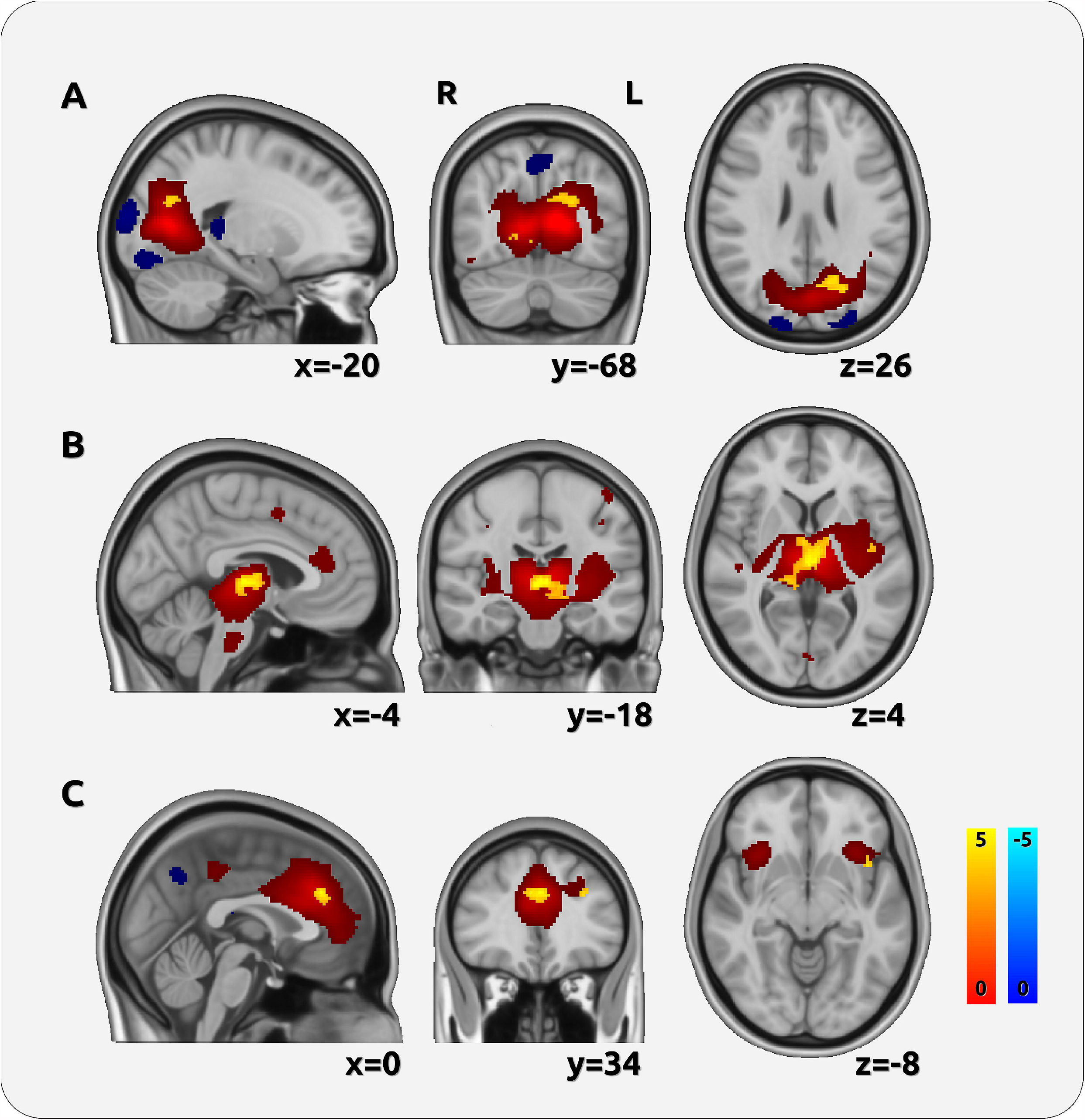
Statistical analysis across the migraine cycle. The Figure shows 3 exemplary intrinsic networks (visual-sensory network, thalamo-insular network, and the salience network). The dark red and blue colours represent the entire extension of the networks. The results of the change of network connectivity throughout the migraine cycle are superimposed in lighter colours. For network 5 (A), we found a main effect in the primary visual cortex in the calcarine sulcus; for network 27 (B), we found an effect predominantly in the thalamus; and for network 34 (C), we observed an effect in the anterior insular cortex and the dorsal ACC. R = right hemisphere, L = left hemisphere.

### Trajectory two

One limbic network followed trajectory two: the connectivity of the parahippocampal gyrus linearly decreased towards the next attack and reached its lowest magnitude during the headache (Table 2c). We did not observe any significant effect of the opposite contrast (linear increases over the pain-free interval with peak magnitude during the headache). All statistical tests were thresholded at p<0.05 (FDR corrected).

## Discussion

The present longitudinal fMRI study aimed to study the cyclic connectivity of intrinsic functional networks in episodic migraine. We analysed two possible network trajectories over the migraine interval.

### Sensory networks

Alterations of sensory processing in migraine have been shown in various studies ^9,14^. Sensory thresholds have been suggested to depend on the phase within the migraine cycle, where the lowest thresholds were detected during the headache phase ^9,10^. Here, we mainly observed linearly increasing sensory network connectivity over the pain-free interval with a peak prior to the beginning of the headache that “dropped” to the initial level during the headache in brain regions belonging to visual (e.g. intracalcarine cortex and occipital pole), auditory (e.g. planum temporale), and somatosensory networks (e.g. postcentral gyrus). For a visual network we also observed the inverse effect, where the connectivity of the lateral occipital cortex decreased over the pain-free interval and reached its peak during the headache. An increasing magnitude of network connectivity in sensory networks over the migraine cycle towards the attack, as observed in our study, is likely to reflect an increased sensitivity to sensory input. Additionally, it may reflect an increasing vulnerability to internal and external stressful events during the pain-free interval. External triggers for migraine attacks in some patients, such as bright or flickering light ^15^ and odours ^16^, highlight dysfunctional sensory processing. The origin of such hypersensitivity in migraineurs is not yet understood. Genetic predispositions, such as ion channelopathies, can affect excitability thresholds making the cortex more vulnerable to diencephalic (thalamus, hypothalamus) and pontine inflow ^17^. Consequently, repetitive stimulation may result in an overload of cortical sensory neurons in migraine. The “drop” of network connectivity during the headache suggests a “rebooting” of cortical processes.

### Salience networks

In line with our hypotheses, we found cyclic changes in the salience network, e.g. in the anterior cingulate cortex and the anterior insula. Similar to the sensory networks, connectivity increased over the interictal interval, reached the maximum prior to the headache, and “dropped” to the baseline level during the attack. These findings are corroborated by previous work on ictal and pre-ictal alterations of the salience network in migraine ^2^.

It could be speculated that the deviances in the salience network are associated with an interictally-impaired habituation in migraine, which could be explained either by reduced intra-cortical inhibition or by increased cortical excitability. Indeed, some studies report an abnormal level of both inhibitory and excitatory neurotransmitters, such as in GABA (gamma-aminobutyric acid) and glutamate ^18^. Specifically, an increased amount of glutamate has been observed in migraine patients in the anterior paracingulate cortex, a region that is an essential part of the salience network ^19^. Numerous neurophysiological studies confirm the impaired inhibition by showing increased responsiveness to sensory stimulation in the visual, auditory, and somatosensory domain (for a comprehensive review see ^14^).

### Limbic networks

Altered limbic connectivity is likely to be responsible for several migraine symptoms during the prodromal, ictal and postictal phases of the migraine cycle (e.g. fatigue, irritability, yawning, polyuria, and food cravings). Furthermore, many trigger factors for migraine attacks are associated with limbic circuits, e.g. psycho-physical distress and emotions, homeostatic changes or circadian rhythms ^20^.

Here, we revealed connectivity changes of the right DLPFC. This region is integrated in a limbic network, which also includes the hippocampus, thalamus, hypothalamus, cingulate cortex, and the insula. The DLPFC connectivity increased over the pain-free interval, reached its peak immediately prior to the headache, and “dropped” to the lowest level during the headache. The DLPFC is a key node of cognitive circuits and contributes to executive function, attention, decision-making, and emotional regulation^21^. The DLPFC further contributes to the top-down control of nociceptive input ^22^.Increasing connectivity of the DLPFC over the pain-free interval may reflect the increasing prefrontal effort to control cognitive, emotional, and sensory processes, whereas a “drop” in DLPFC connectivity during the ictal phase is likely to reflect the altered pain and cognitive processes resulting in headache and cognitive deficits.

In a further limbic network, the connectivity of the left hippocampus (as part of a bilateral hippocampus-amygdala network) increased interictally and “dropped” during the headache. There is growing evidence showing a cortical effect of the hippocampus in migraineurs in comparison to healthy controls, e.g. a reduced volume ^23^ and a dependency of its connectivity on the attack frequency ^24^. The amygdala-hippocampus network has been associated with the processing of mood ^25^, memory encoding ^26^, and emotion-related cognitive functions ^27^. Our findings are in line with these recent studies and could reflect an increase in anxiety, decrease in mood, and pain-related rumination towards the next attack. Indeed, a previous study reported growing connectivity between the amygdala and the hippocampus after stress ^28^.

In addition, a limbic network, composed of the bilateral parahippocampal gyri, hippocampi and the amygdalae, showed a linearly decreasing magnitude of connectivity of the left parahippocampal gyrus over the pain-free interval and reached its lowest level during the headache. The network that comprises these three regions has been associated predominantly with the processing of memory ^29^ and object novelty ^30^. Further, lesions in the parahippocampal gyrus were found to cause severe memory deficits ^31^ and impaired emotional responses ^32^.

The migraineurs’ imperative focus on the headache is suggested to explain the low connectivity of the parahippocampus during the attack. This focus may impair cognitive functions *per se*, but can also prevent the processing of any past- or future-related memory during attacks ^33^.

Contrary to our hypothesis, we did not find any cyclic alterations of hypothalamic networks. One explanation might be that the hypothalamus is not a permanent part of the intrinsic networks, but may instead orchestrate different networks at different time points during the migraine interval as a rhythm generator.

### Thalamo-insular network

There is increasing neurophysiological and neuroimaging evidence for the role of the thalamus within the migraine cycle ^34^. As the thalamus is connected to the sensory, salience, and limbic networks in an upstream direction, almost every stimulus that reaches the cortex is first processed and gated by the thalamus ^35^. We observed bilateral connectivity changes of the thalamus within a thalamo-insular network. Thalamic connectivity changes followed the trajectory of a linear increase over the interictal interval with a peak immediately prior to the headache and a “drop” to the baseline level during the headache. Structural ^36^, functional ^34^, and neurochemical thalamic alterations ^37^ in migraine have been highlighted in recent imaging studies. During spontaneous attacks, altered functional connectivities between the thalamus and cortical regions contributing to pain processing and modulation (e.g. the insula and the orbitofrontal cortex) have been observed ^38^. Furthermore, using diffusion tensor MRI it has been shown that migraine patients exhibited a higher fractional anisotropy in the thalamus during the interictal interval, which normalised during the attack ^13^.

Through the connections between the thalamus and the principal structures of the salience network (anterior insula, anterior cingulate cortex), the current results may suggest a maladaptive gating mechanism in the migraineurs’ brain that usually prevents an overload of irrelevant and unnecessarily salient information to higher cortical areas. In migraine, internal and external input to the cortex is likely to be insufficiently filtered or inhibited by GABAerge thalamo-cortical and cortico-thalamic circuits ^10^.

### Other networks

We also found cyclic connectivity changes in various other cortical (temporal and frontal networks, as well as in cortically distributed networks) and subcortical networks (cerebellar and basal ganglia). Most of these networks followed the trajectory of a linear increase over the interictal interval with a peak prior to the headache and a “drop” to the baseline level during the headache. This applies to the posterior DMN, which has previously been shown to be altered in migraineurs ^4^ and other chronic pain states ^39^. DMN dysfunctions have been related to maladaptive stress responses, which seems to characterise chronic pain patients in general ^39^. Similarly, we found cyclic changes of the central executive network. Although cognitive symptoms are not included as core symptoms to diagnose migraine disease, they are frequently reported by patients during the premonitory, ictal, and postdrome phases ^40^.

## Conclusion

The present results suggest that migraine disease is associated with cyclic alterations of various functional networks, underlining the complexity of the clinical picture. Importantly, network changes evolve early during the migraine cycle, long before the headache is initiated. Therefore, our findings provide further evidence for the use of psychological approaches during the pain-free migraine interval. Cognitive-behavioural interventions, relaxation techniques, or biofeedback, which are aimed at normalising network functions, help patients to regulate major trigger factors of migraine attacks (e.g. psycho-physical stress, sleep-awake rhythms). By modulating sympathetic activity or opioidergic activity, these interventions are likely to prevent a recurring loss of network synchronicity over the migraine cycle.

## Acknowledgements

We thank Dr Stephanie Irving for providing comments and copy-editing the manuscript. We thank Prof Dr Andreas Straube and Dr Daniel Keeser for their comments.

## Key Findings

- Cortical processes of migraineurs undergo cyclic changes, already detectable in the pain-free phase.
- The increasing magnitude of connectivity towards the next attack in several networks reflects the increasing effort to keep the network integrity intact.
- Network decoupling in the ictal phase can explain the variety of ictal and pre-ictal migraine symptoms.
- These findings provide further evidence for the need of early therapeutic approaches during the pain-free interval.

